# An optimized method to infer relatedness up to the 5^th^ degree from low coverage ancient human genomes

**DOI:** 10.1101/2022.02.11.480116

**Authors:** Emil Nyerki, Tibor Kalmár, Oszkár Schütz, Rui M. Lima, Endre Neparáczki, Tibor Török, Zoltán Maróti

## Abstract

**Background:** Current state of art kinship analysis is capable to infer relatedness up to the 5-6th degree from deeply sequenced DNA if the proper reference population is known. Low coverage, partially genotyped, degraded archaic (or forensic) DNA and often unavailable or unknown reference population poses additional challenges, hence kinship analysis from low coverage archaic sequences so far has been possible up to the second degree with large uncertainties.

**Results:** We performed extensive simulations to identify and correct the main factors of bias in kinship analysis from low coverage data. As a result, we introduce a new metric for correction and offer a guideline, which overcomes the difficulties associated with low coverage samples. We validated our methodology on experimental modern and archaic data with widely different genome coverages (0.12x-11.9x) using samples with known family relations and known or unknown population structure. Out of 2526 ancient individuals from the REICH data set we confirmed all 96 indicated, and identified 303 new relatives additionally.

**Conclusion:** With the proposed workflow we provide the necessary additional tools to calculate the corrected kinship coefficient from the commonly used genome data formats. Our methodology allows to reliably identify relatedness up to the 4-5th degree from variable/low coverage archaic (or badly degraded forensic) WGS genome data.

## Introduction

Kinship analysis is a method for determining the familial relationship between individuals from genome data. The kinship coefficient, defined as the probability that two homologous alleles drawn from each of two individuals are identical by descent (IBD), is a classic measurement of relatedness [1], [2]. Several algorithms had been developed such as GERMLINE [3], fastIBD [4], GRAB [5] and ANGSD [6], based on different strategies and metrics of IBD segments for calculating relatedness from whole-genome sequence (WGS) data. Distinguishing IBD (familiar relatedness) from identity-by-state (IBS – due to population relatedness) is difficult as both result in genetic similarity through the sharing of alleles. In spite of the biologically driven variation in IBD sharing - the outcome of the stochastic nature of recombination and segregation - it is possible to infer kinship up to the 5-6th degree of relatedness from deeply typed WGS data. Achieving such a deep level of certainty requires 1, deeply sequenced diploid WGS data, 2, an appropriate set of reference data, and 3, clever algorithms to account for biological variations resulting from familial (IBD) and population structure (IBS).

Recently huge set of genome data have been generated from archaic samples in order to uncover the genetic relations of ancient and modern populations. From these data it is also of high interest to study the family organization of archaic populations, however, analyzing ancient DNA (aDNA) poses additional difficulties due to the widely different but generally low genome coverage and postmortem damage (PMD). The aDNA databases usually contain sequence data between 0.05-3x average genome coverage [7]–[9], since the sequencing of archaic samples with low endogenous DNA-content is still challenging and costly. Differences in coverage and only partial overlap of genetic markers between samples can lead to significant bias when comparing the frequencies and genotype likelihoods of genetic variants, leading to uncertainties of the inferred genotype probabilities. To overcome typing bias, random sampling of one allele per site (pseudo-haploid calling) was used successfully in aDNA studies [10]–[19]. For being able to compare diploid and pseudo-haploid data sets, heterozygous alleles of diploid data need to be random pseudo-haploized (RpsH) by randomly choosing one of the heterozygote alleles as if either homozygote REF or homozygote ALT. Although rare alleles can offer significant improvement in some kinship calculation methods when analyzing high quality WGS data, genotype calling from the whole human genome could lead to excessive, variable amount of false positive calls from low coverage, degraded ancient DNA. To minimize this bias, we restricted our analysis to the already known bi-allelic, high frequency and population informative SNPs of the 1240K REICH data set [20]. A further problem in the analysis of archaic data is that in most cases we lack, or have very limited knowledge of the appropriate reference population data to distinguish IBD from IBS.

In the present study we wanted to address the difficulties of low coverage aDNA data and dissect the main factors (and their effects) that influence kinship calculations. To overcome the issue of reference populations we used the PC-Relate algorithm [21] that proposes a principal component based model-free approach for estimating commonly used measures of recent genetic relatedness, such as kinship coefficients and IBD sharing probabilities, in the presence of unspecified population structure. We applied a combination of techniques to mitigate the genotyping uncertainties and tested their effects and limitations on kinship analysis of low coverage archaic sequences. We used simulation to down sample fully genotyped real NGS data to examine the effect of partial marker overlap between samples and we also explored the effect of reference population choice on the kinship coefficient calculation. Based on our results we came up with a new computational approach which can reliably correct the calculation of kinship coefficient even from poorly genotyped data.

Here we offer a guideline and the necessary additional bioinformatics tools to calculate the corrected kinship coefficient, which overcome the technical limitations; the generally low genome coverage, the postmortem damage, the genotyping uncertainties, the partial overlapping of genetic markers between samples and the selection of reference population. As a proof of concept, we validated our proposed methodology on both experimental modern and ancient data with widely different genome coverages, using samples with known family relations and known or unknown population structure.

## Results

### The effect of random pseudo-haploidization on PCA calculations

Since the selected kinship methodology, PC-Relate, applies principal component analysis (PCA) to identify population structure, first we tested the effect of the random RPsH on the outcome of PCA analysis. According to our results the RPsH does not alter the PCA calculations significantly (Supplementary Figure 1.).

### The effect of random pseudo-haploidization on kinship coefficient calculation

Next, we assessed the effect of RPsH on kinship calculation by selecting four populations with different population structure from the 1000 Human Genome Project Phase III (1KG phase 3) data set [22]. The four populations were as follows: Finnish (FIN), British (GBR), Toscani (TSI), and Utah residents (CEPH) with Northern and Western European ancestry (CEU). In our experiments we generated 6 different pseudo-haploid data sets from the original diploid data using different random seeds.

To study solely the effect of RPsH on kinship coefficient calculation, without the potential interference of other effects, like biological variation (skewed recombination/segregation), differences in sequence alignment, genotyping, genome composition or in the population structure between the relatives, we calculated the “kinship” of the same sample from different randomizations in each case. This idealized experimental setup is equal to the monzygotic (so-called identical) twins kinship relation (or sample matching in forensic) while the maximal expected kinship coefficient (0.5) allows the most sensitive analysis. Since this setup does not exclude differences between the genome structure of the test sample and the reference population we selected one random individual from each of the four populations, with the exception of the admixed CEU population, where we selected 2 individuals (1 male, 1 female) (Table 1).

**Table 1.**
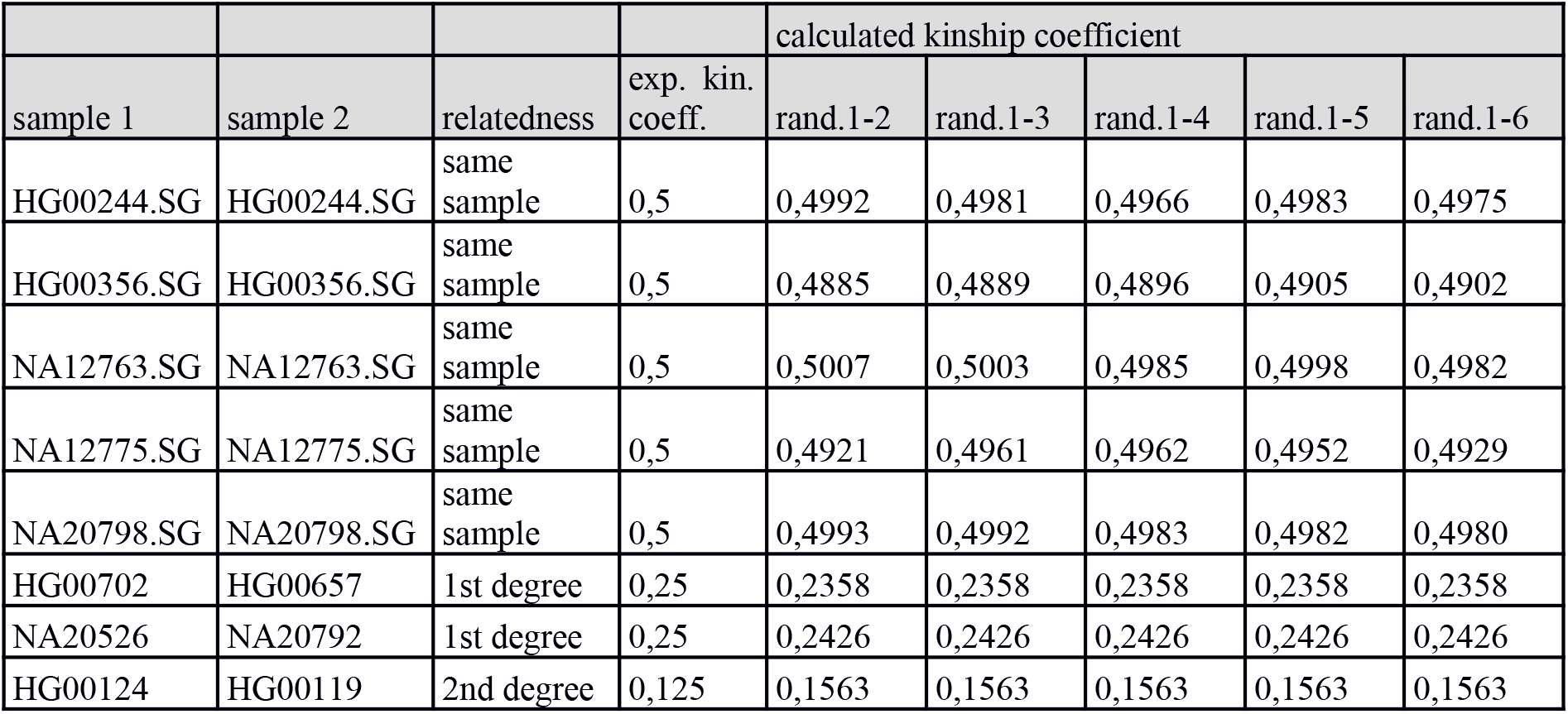
Effect of random pseudo-haploidiziation on calculated kinship coefficient. Each sample was analyzed against its own reference population.

| sample 1 | sample 2 | relatedness | exp. kin. coeff. | calculated kinship coefficient |  |  |  |  |
| --- | --- | --- | --- | --- | --- | --- | --- | --- |
|  |  |  |  | rand.1-2 | rand.1-3 | rand.1-4 | rand.1-5 | rand.1-6 |
| HG00244.SG | HG00244.SG | same sample | 0,5 | 0,4992 | 0,4981 | 0,4966 | 0,4983 | 0,4975 |
| HG00356.SG | HG00356.SG | same sample | 0,5 | 0,4885 | 0,4889 | 0,4896 | 0,4905 | 0,4902 |
| NA12763.SG | NA12763.SG | same sample | 0,5 | 0,5007 | 0,5003 | 0,4985 | 0,4998 | 0,4982 |
| NA12775.SG | NA12775.SG | same sample | 0,5 | 0,4921 | 0,4961 | 0,4962 | 0,4952 | 0,4929 |
| NA20798.SG | NA20798.SG | same sample | 0,5 | 0,4993 | 0,4992 | 0,4983 | 0,4982 | 0,4980 |
| HG00702 | HG00657 | 1st degree | 0,25 | 0,2358 | 0,2358 | 0,2358 | 0,2358 | 0,2358 |
| NA20526 | NA20792 | 1st degree | 0,25 | 0,2426 | 0,2426 | 0,2426 | 0,2426 | 0,2426 |
| HG00124 | HG00119 | 2nd degree | 0,125 | 0,1563 | 0,1563 | 0,1563 | 0,1563 | 0,1563 |

To study the effect of RPsH on true first/second order relatives we also selected samples with known family relations from the 1KG phase3 data set. a) HG00702-HG00657 a parent-child relation from a Han population, where exactly 50% of genome is shared between the two samples, b) NA20526-NA20792 siblings from a TSI population, where 50-50% of the genome comes from the same parents, however a different subset of the markers are found in the sibs due to segregation and recombination, and c) HG00124-HG00119 second order relatives from a GBR population, where statistically 25% of genomes are shared. Then, we calculated the kinship coefficient for each of the selected relatives (kin1/kin2) comparing the data of the first relative (kin1) from the first randomized data set with the data of its relative (kin2) from the other five randomizations using the samples’ own reference populations. Knowing the expected kinship coefficients we could validate that RPsH does not significantly alter the calculated kinship coefficient in these settings (Table 1).

### The effect of overlapping marker fraction on the kinship coefficient calculation

Since kinship coefficient calculation is based on the IBD segments that are shared between two samples, that can be only assessed at points where both samples are genotyped, we defined a metric called overlapping marker fraction. We calculate this metric by dividing the number of markers where both samples are genotyped with the total number of markers in the data set (1240K). To study exclusively the effect of overlapping marker fraction on kinship coefficient calculation we used the “monozygotic twins” scenario with different haploidizations of the same sample as described previously. We performed a stepwise series of marker depletion between each sample pairs in the range between 100% to 5%, by randomly removing markers (genotyped in both samples) in 5% steps. Then calculated the kinship coefficient between the depleted samples using the EUR superpopulation as reference. We repeated this analysis 5 times, with 5 independently randomized datasets, and obtained the same results in all cases. Moreover, the analysis was performed for 5 randomly selected European individuals (Fig.1). The results confirmed that randomization does not affect the outcome, and also revealed that the kinship coefficient is a linear function of the marker overlap fraction, enabling a simple correction of the kinship coefficient value for low coverage genomes.

**Figure 1.**
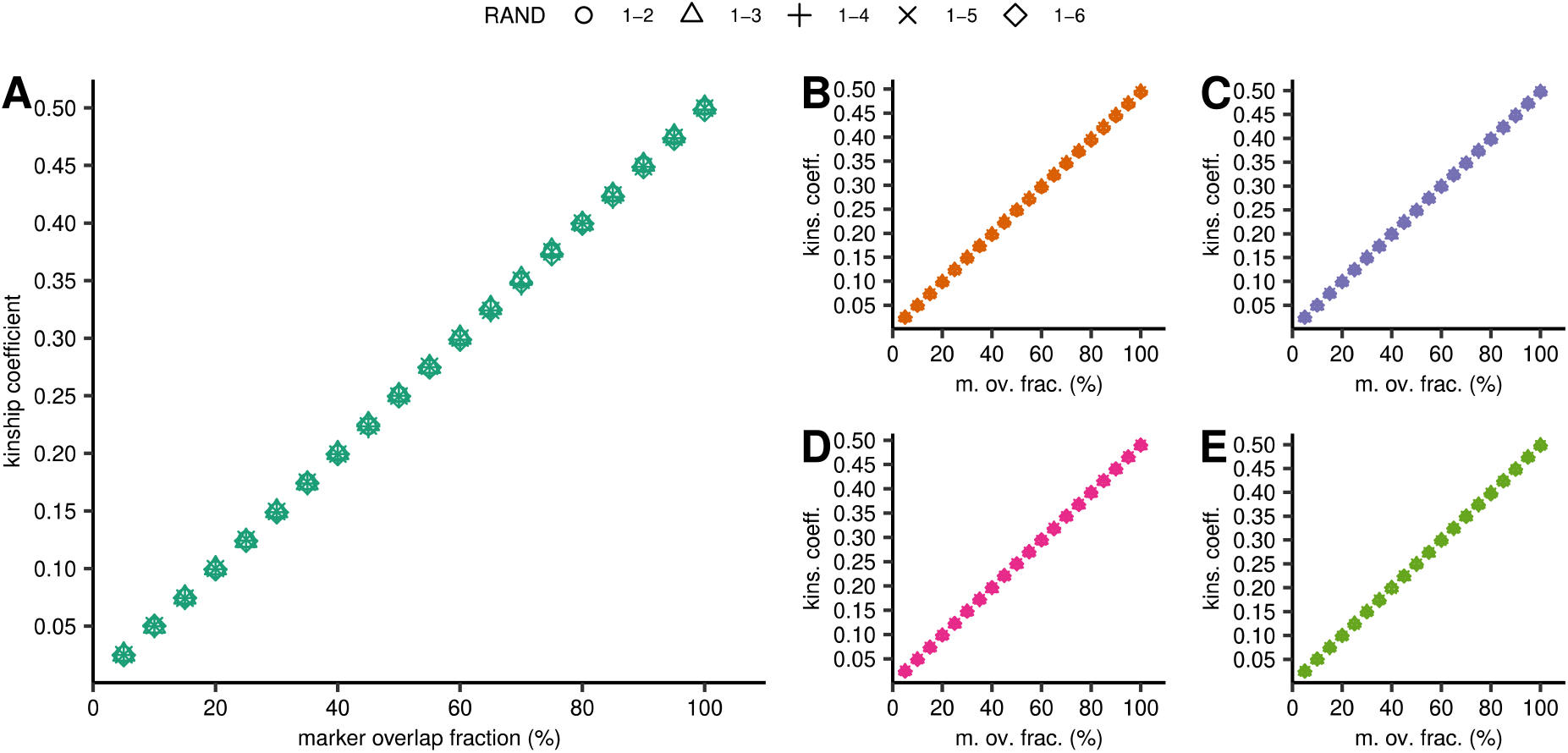
Effect of marker overlap fraction and randomization on the calculated kinship coeficient. In each graph we display the results of five parallel pairwise kinship calculations between the first and the other five randomized data set, in each parallel we calculated the kinship coefficients between different randomizations of the same sample at different marker overlap fractions. The analysis was repeated for 5 European individuals each representing different population structure: A: CEU-female, B: CEU-male, C: GBR-male, D: FIN-female, E: TSI-male). In all cases we used the EUR super population as reference.

### The effect of reference population selection on kinship analysis

Using the same public 1KG phase3 data set and the three known relatives HG00702-HG00657 parent-child of Han population, NA20526-NA20792 siblings of TSI population and HG00124-HG00119 second order relatives of GBR population, we investigated the effect of the reference population on the calculated kinship coefficients. We tested three different scenarios: 1) the reference population was the same as that to which the selected individual belonged to; 2) the reference population was from a different super-population (AFR) than that of the selected individual’s 3) the reference population was from the same super-population (JPT, IBS and FIN) as the selected individual was derived from (Figure 2, Supplementary Figure 2). To study the effect of overlapping marker fractions in these more complex cases, the selected sample pairs were also marker depleted in the range of 100% to 5% overlap fractions. The results revealed that a reference population with significantly different genetic background, like African for European samples, strongly corrupts the results, while a superpopulation with similar genetic background to the sample’s own population gives very similar results as the own population.

**Figure 2.**
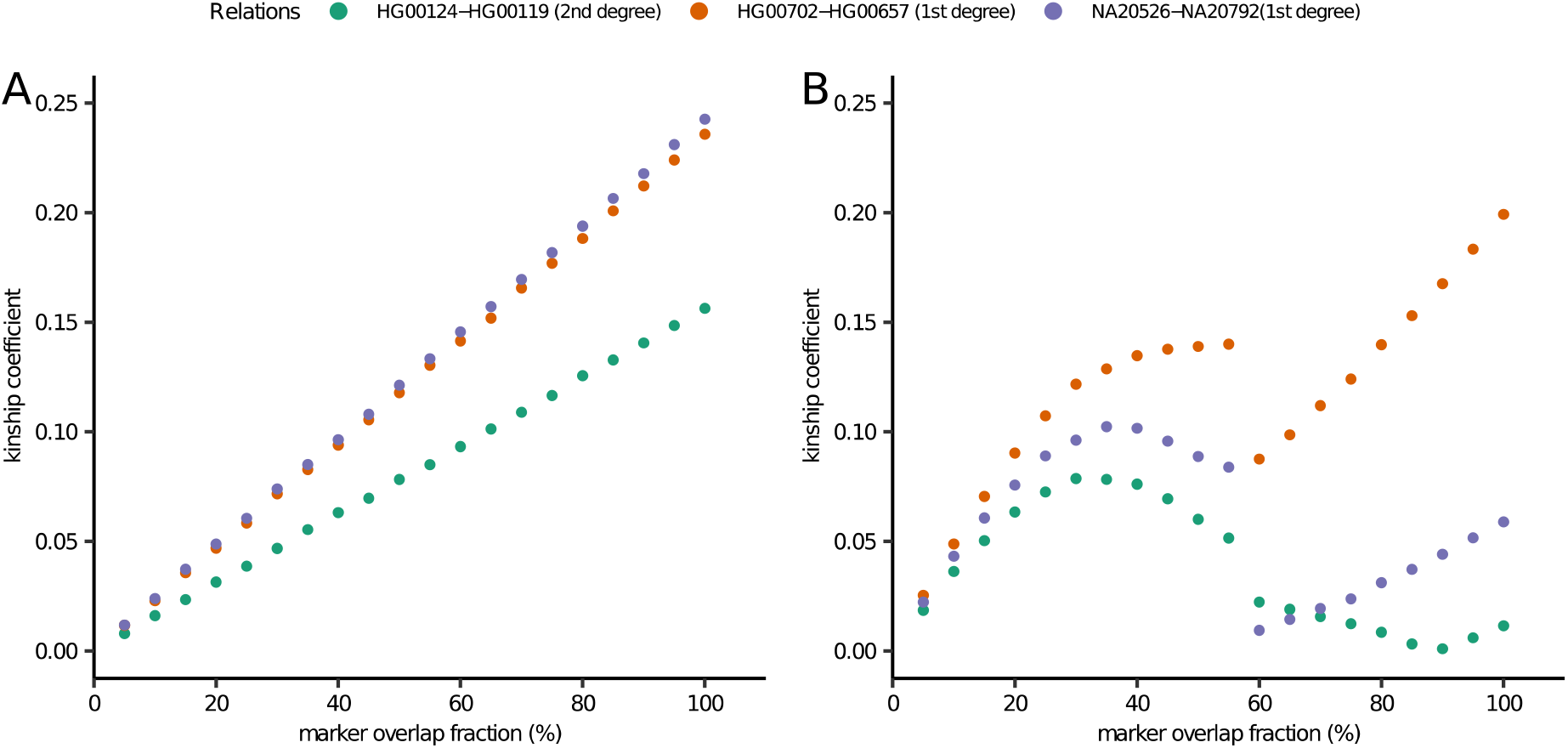
The effect of reference population choice and marker overlap fraction on the calculated kinship coefficients between selected 1^st^-2^nd^ degree relatives. Markers were depleted between the relatives to 5-100% overlap fractions. A: The samples’ own populations used as references B: the AFR super population was used for each sample.

### Effect of reference population selection on kinship analysis in a complex admixed family with multiple ethnic relations

To assess the choice of reference population in the kinship analysis of admixed individuals (often the case in archaic populations) we analyzed a complex, multiply admixed Cabo Verdean-Hungarian family with known pedigree. In this pedigree we had WGS data of siblings (first order), differently admixed half-sibs (2^nd^ order) and 5^th^ order relatives as shown in Figure 3.

**Figure 3.**
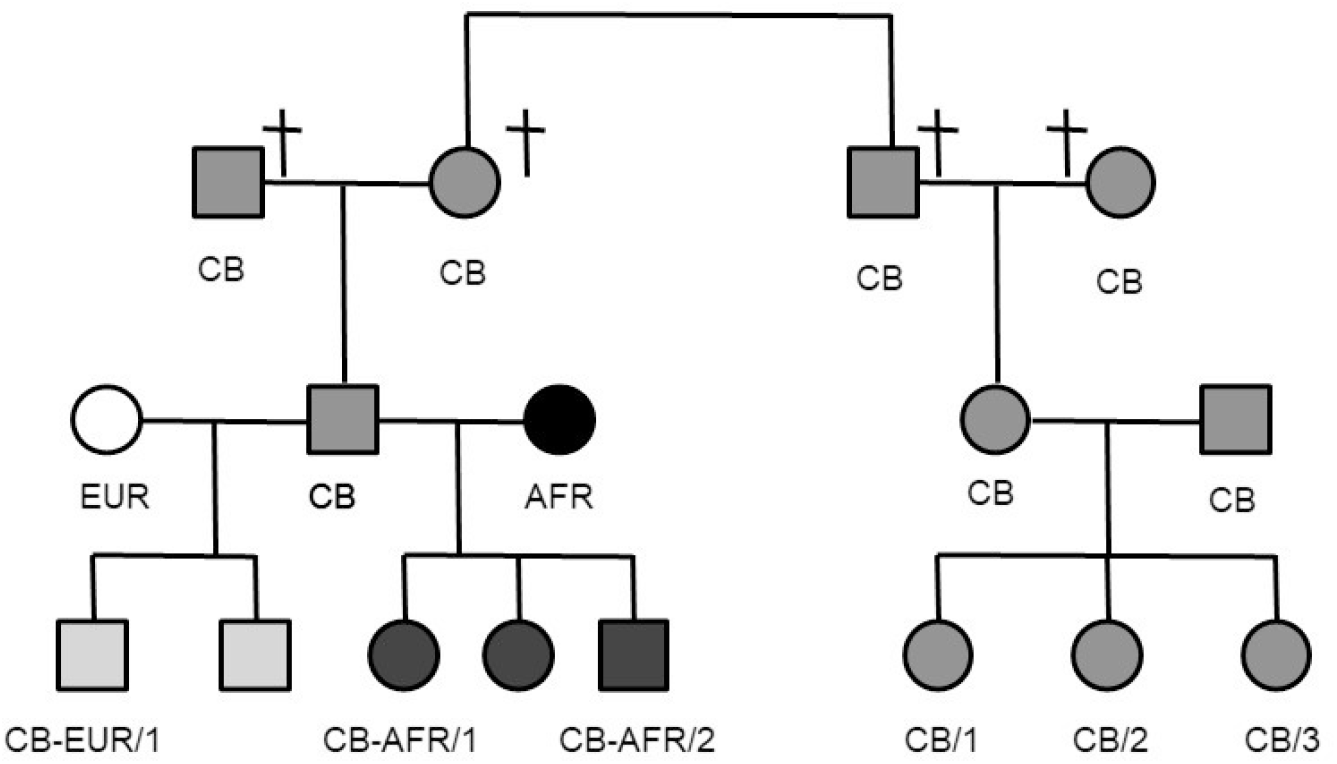
Pedigree of a complex admixed modern family with Cabo-Verdean (CB, 50%-50% old EUR/AFR admix), AFR (100% African), EUR (100% Hungarian) family members and recently admixed offsprings. WGS data was only available from third generation individuals denoted with numbers in their ID.

We tested two scenarios: the reference population was 1) only African (AFR) representing the majority of the admix sources in the tested samples; 2) both African and European (EUR) populations were included as references. Additionally, we performed marker depletion to investigate the effect of coverage in this complex scenario (Figure 4).

**Figure 4.**
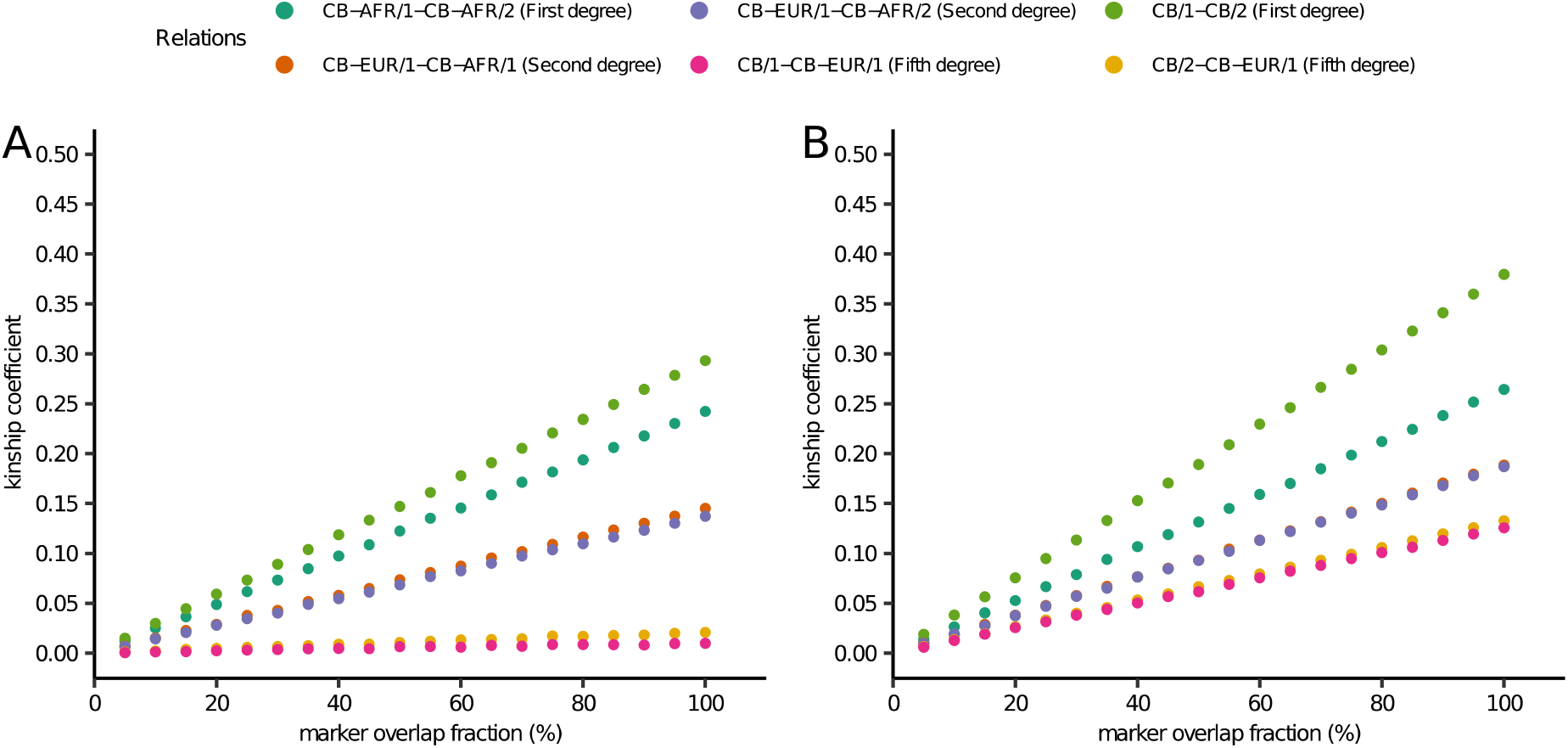
The effect of reference population choice and marker overlap fraction on the calculated kinship coefficient in a complex admixed modern family with 1^st^ (sibs), 2^nd^ (half-sibs) and 5^th^ degree relations, where individuals originated from populations with largely different genetic structure. Markers were depleted between the relatives to 5-100% overlap fractions. A) The reference population was a combined set of AFR+EUR populations, B) The reference populations were AFR populations.

To test whether random haploidization alters the kinship coefficient calculation compared to the better phased diploid data in this complex admixed case we also performed this analysis from the original diploid data set (Supplementary Table 1.) with the AFR + EUR reference population. The results in Figure 4 demonstrate, that in order to obtain realistic coefficient values in case of a complex admixture, a combined set of reference populations is required, representing the population structure of all ancestors. Using just the majority source as reference significantly distorts the result. It was confirmed again, that even in such complex admixed family the differences due to RPsH were negligible.

### Correction of the calculated kinship coefficient

According to the idealistic monozygotic twins scenario the paired marker overlap fraction between the depleted samples had a strong linear correlation with the calculated kinship coefficients offering a simple mathematical correction for the coefficients of partially overlapping data.

The kinship coefficients calculated from the full marker set (100 % marker overlap) resulted in a value close to the expected one deduced from the known degree. Dividing the calculated kinship coefficients by the overlapping marker fractions of the analyzed samples we obtained the corrected kinship coefficients, which are nearly identical to the one with 100% marker overlap, across all simulated genome coverage ranges. (Figure 5.)

**Figure 5.**
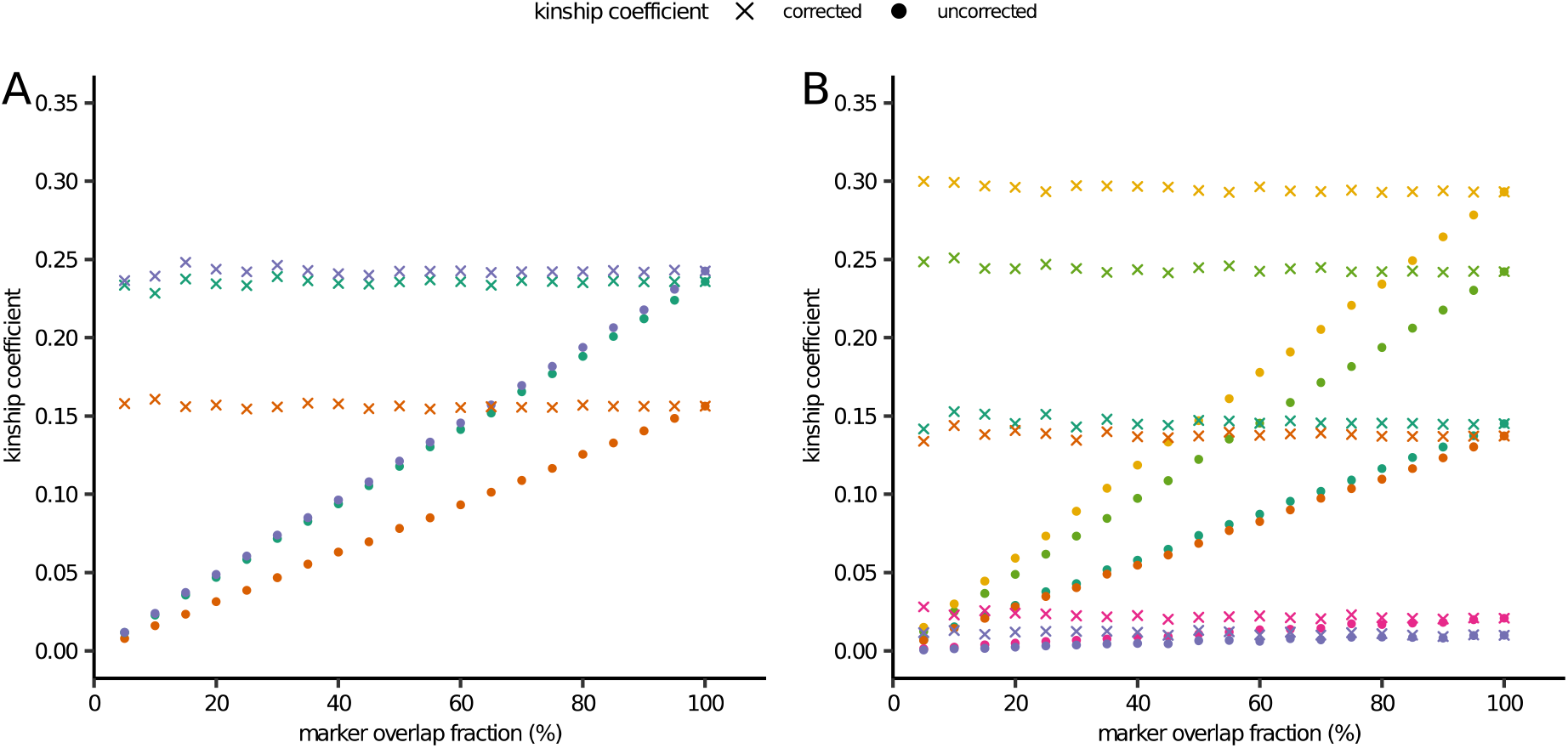
Uncorrected and corrected kinship coefficients relative to the fraction of marker overlap between A) 1KG 1^st^ and 2^nd^ degree relatives, B) in a complex admixed modern family (1^st^, 2^nd^ and 5^th^ degree relatives). Markers were depleted between the relatives to 5-100% overlap fractions. The colors represent kinship coefficients between different known kinship relations. The non-corrected kinship coefficients are displayed as solid dots, the corrected values are shown as X. Each relation was analyzed using the appropriate reference population.

### Kinship analysis of ancient samples with known relations using kinship coefficient correction

To show that our methodology is also suitable for archaic data we analyzed low coverage archaic sequences with known family relations. In the first example we present the analysis of a known father-son (first degree) relation of two Medieval samples. From both remains we had two types of biological samples (bone powder) one taken from the teeth and other from the *pars petrosa*. From the father we had three parallel DNA isolates and NGS libraries, two prepared from pars petrosa and an additional one from teeth. From the offspring we had one DNA isolate and NGS library prepared from both types of biological samples. Altogether we had 3+2 NGS sequences with largely different genome coverages (0.87x-11.9x) from these two archaic individuals. Accordingly, we assessed the robustness of our correction method on the 6 combinations of these data sets. We present the uncorrected and corrected kinship coefficients calculated from these data in Table 2.

**Table 2.** Correction of kinship coefficient for different genome coverage of archaic samples with known first degree relation. Phrases high, medium and low refer to coverage levels.

| sample 1 | sample 2 | marker overlap fraction | relation | expected kinship coeff | uncorretcted kinship coeff | corrected kinship coeff |
| --- | --- | --- | --- | --- | --- | --- |
| father high | child high | 0.93378 | first degree | 0.25 | 0.22883 | 0.24506 |
| father high | child low | 0.37760 | first degree | 0.25 | 0.09179 | 0.24308 |
| father medium | child high | 0.73008 | first degree | 0.25 | 0.17604 | 0.24112 |
| father medium | child low | 0.29601 | first degree | 0.25 | 0.07150 | 0.24154 |
| father low | child high | 0.51298 | first degree | 0.25 | 0.12239 | 0.23858 |
| father low | child low | 0.37806 | first degree | 0.25 | 0.05014 | 0.24095 |
| father high | father medium | 0.77832 | sample matching | 0.5 | 0.38284 | 0.49189 |
| father high | father low | 0.54719 | sample matching | 0.5 | 0.27020 | 0.49379 |
| father medium | father low | 0.42839 | sample matching | 0.5 | 0.20863 | 0.48700 |
| child high | child low | 0.35433 | sample matching | 0.5 | 0.17286 | 0.48785 |
| sample | sample type | coverage | typed marker count (1240k) |  |  |  |
| father high | teeth | 11.997 | 1148973 |  |  |  |
| father mid | petrosa | 3.057 | 896623 |  |  |  |
| father low | petrosa | 1.546 | 630405 |  |  |  |
| child high | petrosa | 5.510 | 1075845 |  |  |  |
| child low | petrosa | 0.879 | 435005 |  |  |  |

In the second example we reanalyzed a published group of five related males from the Corded Ware Culture (2500-2050 BCE) with first, second, third and fourth degree kinship relations [16]. In Table 3 we present the family relations with uncorrected and corrected kinship coefficients calculated by our methodology.

**Table 3.**
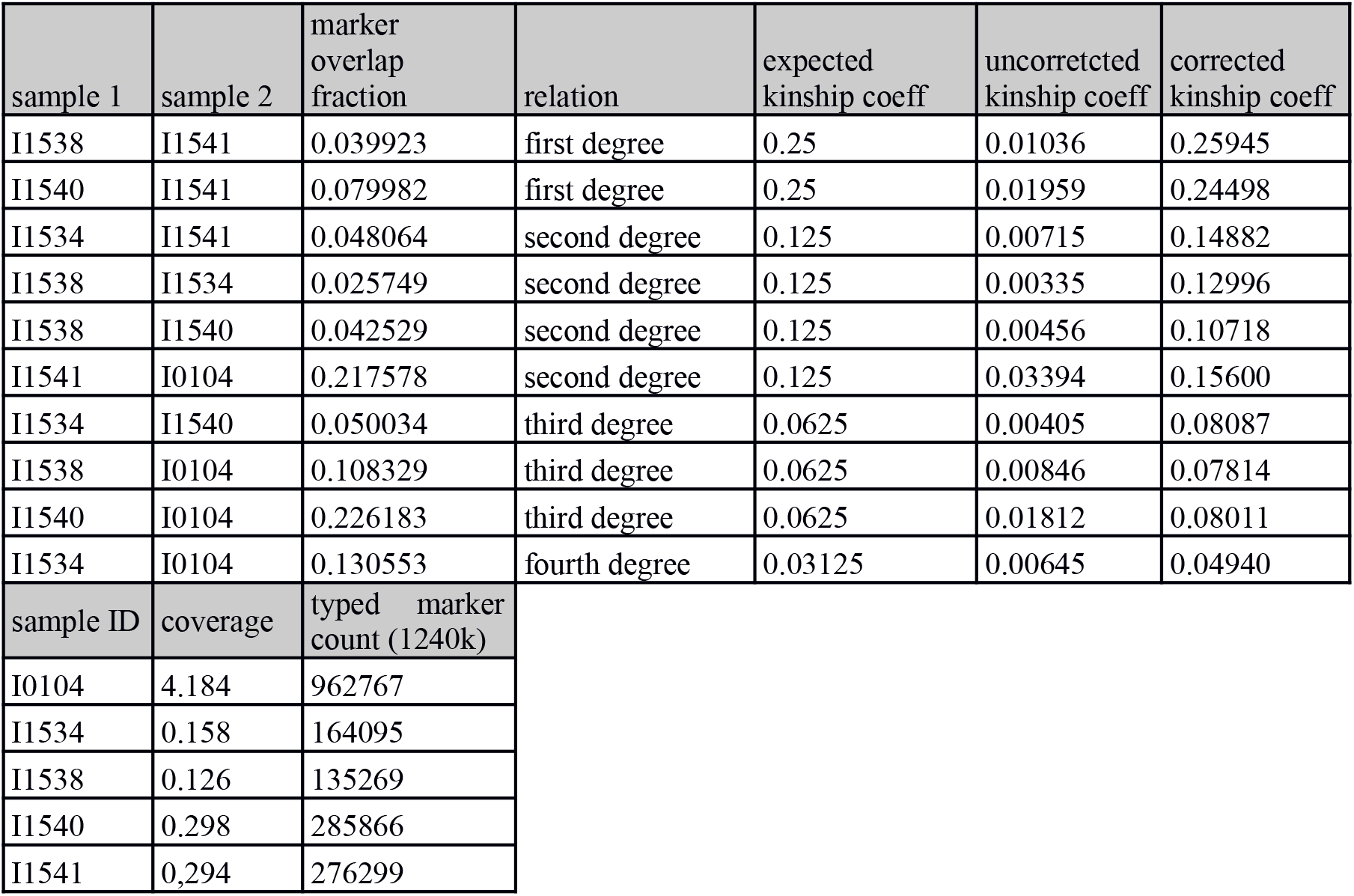
Kinship analysis of a large archaic Corded Ware family with multiple 1st to 4th degree of relations. These individuals were analyzed in the original READ manuscript where relations were identified up to the 2^nd^ degree except the one between I1538 and I1540, and all of the 3^rd^ and 4^th^ relations were inferred from the family relations.

### Kinship analysis of ancient samples from the REICH 1240K data set

We performed kinship analysis on 2526 ancient individuals from the REICH 1240K data set that had more than 100k genotyped markers. Since no manual curation or additional matching reference population was used we filtered kinship relations only up to the 3rd degree. Our analysis identified all sample duplicates, joint datasets of the same sample and all but one of the 96 previously identified kinship relations (Supplementary Table 2.). In case of I8503, an indicated possible 1st degree relative of I8531 there was only 1.6% marker overlap and this relation was only excluded because of the applied stricter threshold. Manual filtering of the sample gave 0.003308 uncorrected, and 0.198185 corrected kinship coefficient confirming the first degree relation even in these very low coverage sample pairs. In addition to the published data, our approach indicated 303 new relatives (Supplementary Table 2). In a handful of cases where no proper reference population was present in the data set the correction resulted in invalid distant (3rd - 4th) kinship relations, these sample groups were highlighted in Supplementary Table 2.

## Discussion

Identification of relatives from the genomic data of ancestors on the one hand is of great interest, as it allows the study of family relationships, but it is also a precondition for most population genetic analyses to exclude close relatives from data sets (e.g. ADMIXTURE, PCA) because their inclusion would distort them. Up to now, the best analysis tools were able to indicate only first and second degree relatedness from low coverage ancient samples. For example, a recent heuristic method READ (Relationship Estimation from Ancient DNA) offers to infer relatedness up to 2^nd^ degree from as low as 0.1x coverage sequence data [23]. In the most comprehensive REICH2020 archaic genome data set [20] the majority of the indicated kinship relations are 1^st^ degree and the handful of indicated 2^nd^ degree relations in all cases are uncertain (the samples are labeled with 1d.or.2d.rel tag).

Diploid variant calling and genotype likelihood based methods with the extra information of rare alleles allow better phasing and identification of IBD fragments leading to improved kinship coefficient estimations from deeply genotyped WGS data. However these techniques introduce major bias due to the huge differences in the genotype likelihoods or diploid variant calling from low/different genome coverage aDNA. To overcome these difficulties and mitigate the main genotyping biases in case of low coverage archaic samples we used a combination of strategies to account for the effects caused by PMD and varying low genome coverage. We used random allele sampling, that is the gold standard methodology when performing PCA and other population genetic analyses on ancient samples, as it leads to statistically equal genotype likelihoods of genotyped markers regardless of the genome coverage. To avoid excessive, variable amount of false positive variants due to the variable rate of PMD, exogenous DNA contamination and technical errors (alignment artifacts), we restricted our analysis to the already known biallelic, high frequency and population informative SNPs of the 1240K REICH data set. This strategy perfectly aligned with our choice of kinship analysis method since the PC-Relate algorithm uses PCA to differentiate between IBD/IBS fragments.

We have demostrated that pseudo-haploid data in our analysis pipeline does not affect the result of kinship analysis (Table 1). This is also confirmed by the PCA analysis, showing that the same modern individual from diploid or different pseudo-haploidized data had nearly identical PCA components (Supplementary Figure 1). The kinship coefficient between different pseudo-haploid randomization of the same sample always resulted in kinship coefficient of 0.5 as expected in case of sample matching (or monozygotic twins).

Kinship coefficient calculation is based on the shared IBD fragments of two samples, thus the coverage metric of a single sample alone (used in prior protocols) is not a good measure to quantitatively characterize the relations between pairs of samples. For this reason we defined a new metric, called overlapping marker fraction (the number markers that are typed in both samples divided by the total number of markers in the data set) that according to our study is the major factor influencing the calculated kinship coefficient in our analysis pipeline. Our simulations revealed that the overlapping marker fraction and the calculated kinship coefficient had a strong linear correlation (Figure 1).

Although the PC-Relate algorithm does not require the specification of the underlying population structure of the analyzed relatives, we have shown that a proper reference set is required for the analysis. As expected, the samples’ own reference population resulted in proper kinship coefficients, but using reference from a different super-population flawed the results (Figure 2, 4B). On the other hand, using the samples’ super-population as reference resulted comparable, although slightly higher kinship coefficients compared to the proper reference population (Supplementary Figure 2.) proving the robustness of the PC-Relate algorithm. We also tested the effect of reference population choice in a complex Creole/European admixed Cabo Verdean-Hungarian family with known 1^st^ to 5^th^ degree family relations. We have shown that the best result is achieved when all super-populations of the sources are included in the reference population set (Figure 4). Comparing the analyses of pseudo-haploid and diploid data for this complex admixed family confirmed the robustness of our approach, as we got nearly identical results (Supplementary Table 1.).

We simulated partially genotyped samples (associated with low genome coverage) by artificially down sampling the data of fully genotyped real modern samples in a controlled fashion in several experiments. Our simulations revealed that the marker overlap fraction of the samples had the largest effect on kinship coefficient calculation with the PC-Relate based algorithm. The applied correction based on the overlapping marker fractions between poorly genotyped samples, combined with the selection of the appropriate reference population was sufficient to obtain nearly the same kinship coefficients as expected from the pedigree. The applied method worked in both unadmixed and complex admixed pedigrees, which had multiple admixtures and source populations with widely different genome structure. (Figure 5 and Supplementary figure 2-4).

We also confirmed the robustness of our methodology on real archaic data with known family relations. Our analysis showed that in the case of a medieval Hungarian family a general modern European reference super-population gave appropriate results. Despite the fact that the uncorrected kinship coefficients varied highly due to the different genome coverages, our robust methodology resulted in reproducible corrected kinship coefficients consistent with the known family relation in each case (Table 2). In the second example we re-analyzed published kinship relations from the Corded Ware Culture [23]. Compared to the READ software that could indicate relations up to the second degree of kinship, and even missed one second degree relation, our approach could properly identify all relations up to 4^th^ degree from this large ancient family with very low/variable genome coverages (0,12x-4,18x), underlining the efficiency and usefulness of our approach [Table 3].

Our results exposed both the advantages and the limitations of the applied methods and strategies. Although RPsH combined with the choice of the 1240K marker set in our study allowed to overcome genotyping bias of low coverage ancient samples, we have to note, that it restricts the analysis to populations that are properly represented by these markers. PC-Relate uses PCA to differentiate between IBD and IBS, during the analysis more distant ancestry (IBS) is regressed out based on the reference population data. Thus, an improper reference population, or marker sets that are lacking the population specific markers of the tests leads to under estimation of IBS and inflated kinship coefficient estimation. The larger the difference between the structure of related individuals and the reference populations the larger fraction of IBS is accounted incorrectly as IBD, which can seriously bias small kinship coefficients (very distant kinship relations). Accordingly, the 1240K set is less suitable for the analysis of extremely old samples, and for small isolated populations, because these supposedly have less informative markers in this marker set, and also have insufficient reference populations in the current genome databases. The unsupervised analysis of 2526 ancient individuals of the 1240K REICH data set (Supplementary Table 2.) demonstrated that our method can identify real 1^st-^4^th^ degree of relatedness from very low coverage ancient damaged samples and fails only when the proper reference population is not present in the data set. On the other hand, our results show that the 1240K marker set was sufficient to properly analyze 4000 years old ancient Corded Ware Culture individuals with a modern Eurasian reference population, suggesting that the majority of the high frequency EUR informative markers were already fixated at this age. The increasing number of aDNA studies will expectedly identify proper reference populations and suitable high frequency marker sets for still hard to analyze samples.

To facilitate the testing and usage of our approach, we provide a practical workflow (Figure 6, Supplementary Note 1) for kinship analysis of low coverage genome data. Our workflow is based on publicly available free software ANGSD, PLINK and PCAangds. We also provide Kinshiptools [github link] to calculate the pairwise overlapping marker fraction between samples and perform RPsH of diploid data in the commonly used genotype formats, PLINK and EIGENSTRAT.

**Figure 6.**
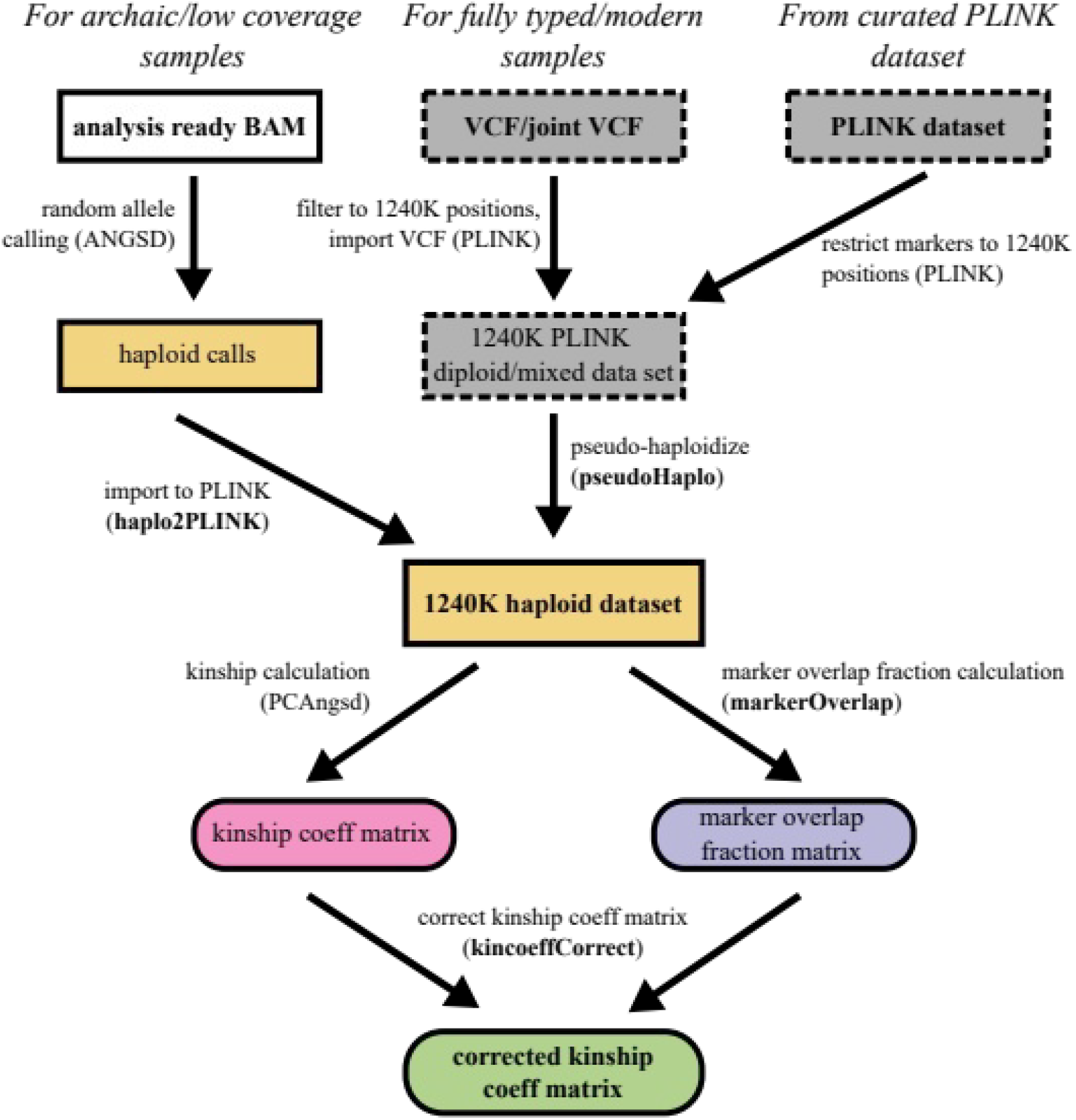
Step-by-step workflow to analyze kinship relation of low coverage archaic/modern samples. Additional tools presented in this manuscript are denoted with bold letters.

In summary our proposed methodology is capable to reliably identify the relatedness up to the 4-5^th^ degree from low coverage genome data, redefining the limits of kinship analysis from low coverage archaic (or badly degraded forensic) WGS data.

## Materials and Methods

### Used data sets

In all of our analysis we used the genome coordinates of 1240K SNP set from Allen Ancient DNA Resource [20]. For marker overlap simulations we used two different full typed modern data sets: the 1000 Human Genome Project Phase 3 data [22]; and a large admixed Cabo Verdean-Hungarian family of known pedigree with first (siblings), second (half siblings, and fifth degree relatives from our anonymized clinical biobank. The variants of the joint VCF (Variant Call Format) files were filtered for the 1240K SNP coordinates and imported into plink 1.9 binary format [24].

Experimental archaic test data was obtained from REICH lab V42.4 1240K data set [20] and our own unpublished archaic sequences. A subset of 2526 ancient individuals from the REICH 1240K were used to test our methodolgy. We filtered ancient samples with >100K genotyped markers. We excluded 6 ancient African individuals and also Mesolithic or older samples because of lacking proper reference populations in the data set. The PLINK binary 1240K genotype data set of our modern and the data set of the ancient CWC individuals described in this manuscript were deposited at XXX [datadeposit link].

### Random pseudo-haploidization and pairwise overlapping marker fraction calculation

We created tools to perform RPsH (pseudHaplo) and to calculate the pairwise marker overlap fraction matrix (markerOverlap) from the main genotype data formats (PLINK and EIGENSTRAT/PACKEDANCESTRYMAP) [git-hub deposit link]. We defined the overlapping marker fraction between two samples as the number of markers typed in both samples divided by the number of all markers in the data set.

Using these tools we created 6 randomly pseudo-haploidized data sets from the fully-typed modern diploid data set using different random seeds. In all of the presented examples we used markerOverlap to calculate the pairwise overlapping marker fraction matrix of samples used for kinship coefficient correction. [Supplementary Note 1].

### Principal component analysis

We selected the EUR super population from 1KG data set (404 samples) and randomized 1-1 samples from each populations (GBR,TSI,IBR,FIN) with different seeds. We performed smartpca [25], [26] analysis on the original data set (489 samples) and used the lsqproject: YES option to project the different randomization of these 50 samples on it. We used the R (version 4.0.5) [27]; and the ggplot2 R package (3.3.5) [28] to visualize our results.

### Simulating the effect of low coverage from fully typed modern data sets

To study the effect of coverage and the resulting lower genotyping percentage on the kinship coefficient calculation in a controlled fashion we created a tool (markerDeplete) to randomly deplete markers from a fully typed (PLINK, EIGENSTRAT) data set resulting in the desired percentage of marker overlap between two samples [github link]. Using this tool we simulated the overlapping marker fraction in the selected samples in the range of 5%-100% with step of 5 percentages.

### Kinship analysis

Kinship analysis was performed by the PCAangsd [29] software (version 0.931) from the ANGSD package [6], that implements a fast parallelized kinship calculation from PLINK or EIGENSTRAT format based on the PC-Relate algorithm [21] with the “*-inbreed 1 -kinship”* parameters. We used the R (version 4.0.5) [27]; the RcppCNPy R package (version 0.2.10) to import the Numpy output files of PCAangsd and the ggplot2 R package (3.3.5) [28] to visualize our results.

## Author Contributions

Conceptualization, methodology, software Z.M. Formal analysis E.Ny., Z.M. Resources E.N., R.L. and T.T. Interpretation of results Z.M., T.K. and E.Ny. Visualization O.S., E. Ny. and Z.M. Writing original draft E.Ny., Z.M. and T.K. All authors took part revising the results and contributed to the final manuscript.

## Acknowledgement

E.Ny. was supported by the ÚNKP-21-3-SZTE-67 New National Excellence Program of the Ministry for Innovation and Technology, from the source of the National Research, Development and Innovation Fund. This research was funded by grants from the National Research, Development and Innovation Office (TUDFO/5157-1/2019-ITM; TKP2020-NKA-23 to E.N.). Rui M. Lima was supported by the NTP-NFTÖ-18 Scholarship.

