## Supplementary Figure 1-2 for "An optimized method to infer relatedness up to the 5^th^ degree from low coverage ancient human genomes"

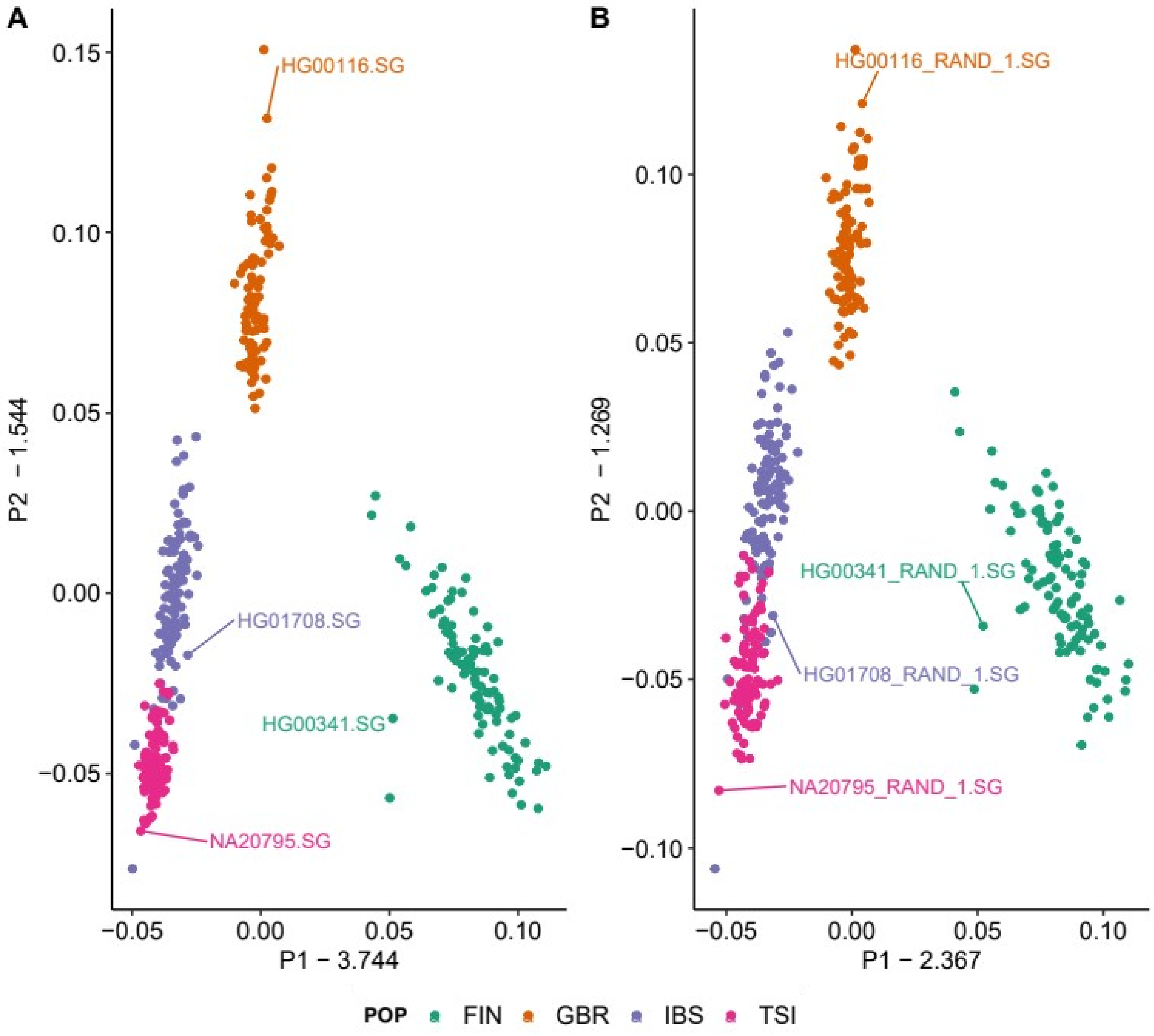


Supplementary Figure 1. Random pseudo-haploidization (RPsH) of diploid data does not significantly alters PCA analysis. A) PCA from the original diploid data set B) the RPsH data set. The EUR super population was used from the 1KG phase 3 data set. We marked one individual from each population on both data set for better illustration.

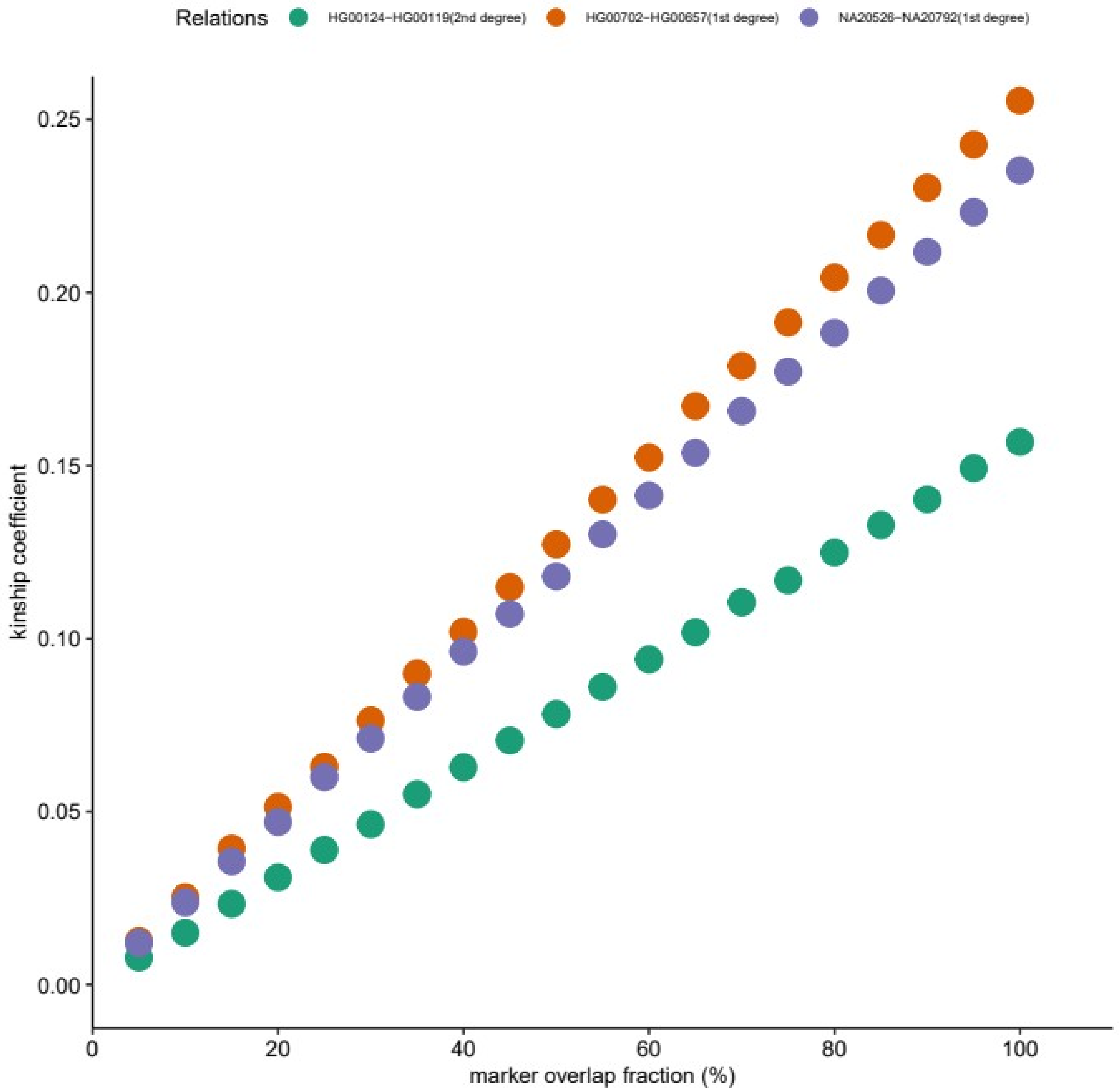


Supplementary Figure 2. The effect of reference population choice and marker overlap fraction on the calculated kinship coefficients between selected $1^{st}$ -$2^{nd}$ degree relatives. Markers were depleted between the relatives to 5-100% overlap fractions. The reference population was from the same super-population (JPT, IBS and FIN) as the selected individual (HAN, TSI and GBR) was derived from.
